# Detection of complex genetic architecture using two-locus population differentiation: modeling epistasis

**DOI:** 10.1101/2020.07.01.182840

**Authors:** Minjun Huang, Britney Graham, Ge Zhang, Jacquelaine Bartlett, Jason H. Moore, Louis Muglia, Scott M. Williams

**Affiliations:** Department of Genetics, Dartmouth College, Geisel School of Medicine, Hanover, NH; Department of Population and Quantitative Health Sciences, Case Western Reserve University, Cleveland, OHIO, USA; Division of Human Genetics, Cincinnati Children’s Hospital Medical Center, USA; The Center for Prevention of Preterm Birth, Perinatal Institute, Cincinnati Children’s Hospital Medical Center, USA; March of Dimes Prematurity Research Center Ohio Collaborative, USA; Department of Pediatrics, University of Cincinnati College of Medicine, USA; Institute for Biomedical Informatics, University of Pennsylvania, Philadelphia, PA; Departments of Population and Quantitative Health Sciences and Genetics and Genome Sciences, Case Western Reserve University, Cleveland, OHIO, USA

**Keywords:** Allelic coevolution, epistasis, preterm birth, population disparity

## Abstract

Recent advances in genetics have increased our understanding of epistasis as important in the genetics of complex phenotypes. However, current analytical methods often cannot detect epistasis, given the multiple testing burden. To address this, we extended our previous method, Evolutionary Triangulation (ET), that uses differences among populations in both disease prevalence and allele frequencies to filter SNPs from association studies to generate novel interaction models. We show that two-locus ET identified several co-evolving gene pairs, where both genes associate with the same disease, and that the number of such pairs is significantly greater than expected by chance. Traits found by two-locus ET included those related to pigmentation and schizophrenia. We then applied two-locus ET to the analysis of preterm birth (PTB) genetics. Using ET to filter SNPs at loci identified by genome-wide association studies (GWAS), we showed that ET derived PTB two-locus models are novel and were not seen when only the index SNPs were used to generate epistatic models. One gene pair, *ADCY5* and *KCNAB1 5’*, was identified as significantly interacting in a model of gestational age (*p* as low as 3 × 10^−3^). Notably, the same ET SNPs in these genes showed significant interactions in three of four cohorts analyzed. The robustness of this gene pair and others, demonstrated that the ET method can be used without prior biological hypotheses based on SNP function to select variants for epistasis testing that could not be identified otherwise. Two-locus ET clearly increased the ability to identify epistasis in complex traits.

---

The frequencies of alleles that affect disease risk or phenotypic variation should parallel the disease or phenotype prevalence distribution among populations. This has recently been shown to be the general rule when comparing GWAS (Genome-Wide Association Studies) associated variants to population background (Guo et al. 2018). A classic example of this pattern is the distribution of rs182549-T, the variant in lactase (*LCT*) that confers lactase persistence (LP). The frequency of this allele is high in populations of northern and western European ancestry that consume cow’s milk into adulthood, but low in populations that do not continue to consume cow’s milk and are mostly not LP (Enattah et al. 2002). Human pigmentation variation also follows this pattern, with alleles that affect skin-color showing extreme divergence among European, African and Asian populations (Sturm 2009). Additionally, SNPs associated with complex traits, such as height and waist-to-hip ratio, are significantly more differentiated across African, Eastern Asian and European populations, compared to non-associating SNPs chosen on the basis of similar minor allele frequencies and linkage disequilibrium scores (Guo et al. 2018). Such parallel patterns for many traits have inspired outlier-detection methods for natural selection and putative disease association (Akey et al. 2002; Oleksyk et al. 2008).

Many patterns of genetic differentiation among populations are due to selective differences that affect allelic distributions. For example, Akey *et al*. identified 174 genes under selection that exhibited extreme interpopulation differentiation compared to the genome-wide background. Of these 174 genes, 17 cause Mendelian diseases or associate with complex diseases (Akey et al. 2002). Unfortunately, most interpopulation comparisons cannot distinguish selection signals from differentiation due to random changes in allele frequencies, leading to high false positive rates with outlier-detection methods (Yi et al. 2010). This could partly explain why only two of the 174 candidate selection genes, *CFTR* and *F5*, have previously been suggested, or confirmed, to be under selection (Akey et al. 2002). That 17 genes do, in fact, associate with disease indicates that allelic population differences are likely important and detectable when there are strong single locus effects. However, for subtler genetic effects, alternative analyses are required to find these patterns (Huang et al. 2016).

Evolutionary Triangulation (ET) (Huang et al. 2016), a method we developed by leveraging patterns of differentiation of SNP allele frequencies across three populations, uses the concept that three-way comparisons can be more effective than pairwise allele frequency comparisons for identifying potential disease associating variants (Figure 1). For example, suppose the prevalence of a specific trait in three populations, A, B and C, is such that population A has a higher prevalence of the trait than either populations B or C, but the prevalences of B and C are similar to each other. Using this pattern of prevalence and population genetic differences, as measured by *F*_ST_, we identified ET SNPs as those that are highly differentiated between A and both B and C, but similar in B and C (Figure 1). We then assessed whether these ET SNPs correlate with traits that are distributed similarly among the three populations, under the assumption that similar patterns of SNP and trait distributions could be used to filter for putatively associating variants. We showed, in several cases, that the ET variants were significantly enriched for genes associating with traits distributed in parallel to the genetic variants. The results were clearest for traits with simple genetic architecture. For traits that have a single locus etiology, i.e. Mendelian disorders, ET was successful in identifying the causative genes for phenotypes, such as lactase persistence, oculocutaneous albinism, and G6PD deficiency, without having to know the phenotypes of individuals whose genotypes were compared. For moderately simple phenotypes, those due to only a few loci, e.g., melanoma, ET was able to identify known risk genes. However, ET did not perform well for traits that have presumably more complex genetic architecture and strong environmental risks, such as Type 2 diabetes mellitus (T2DM). In the case of T2DM, Corona *et al.* did show that the frequencies of 13 known risk alleles decrease with human migration out of Africa, corresponding to the T2DM prevalence among populations (Corona et al. 2013). Similarly, Daub *et al.* showed that specific pathways that are involved in immune response also showed more extreme differentiation than expected due to random processes. They suggested that this was due to adaptation to novel pathogenic variation (Daub et al. 2013).

**Figure 1.**
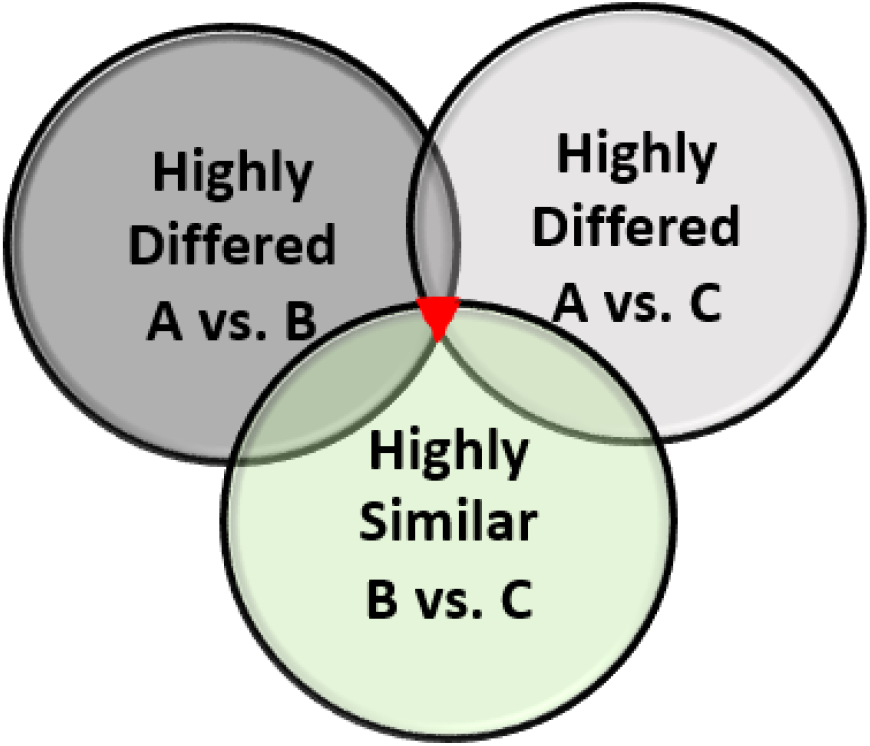
Evolutionary Triangulation (ET) is a method that leverages patterns of genetic differentiation across three populations to filter association results. Suppose the prevalence of a specific trait in three populations, A, B and C is known. Population A has a higher prevalence of the trait than either populations B or C; B and C have similar prevalances of this trait. Using this pattern of divergence, we identified ET SNPs (red triangle), as those that are highly differentiated between A and both B and C, but similar between B and C. By this pattern of overlapping genetic differentiation among population, ET significantly reduces unrelated noises that are not related to our trait of interest.

To improve the detection sensitivity for variants with subtler genetic effects, this paper extends ET to use *F*_ST_ calculated at two loci simultaneously, under the hypothesis that many traits are due to allelic co-variation at two (or more) loci. Unlike the single SNP ET approach, when analyses are performed to detect two-locus coordinated differentiation, we are most likely to identify loci under polygenic selection. A coordinated differentiation pattern of two alleles acting epistatically has been supported by both simulation and empirical analyses. In simulations, Takahasi and Tajima showed in a two-locus system that when the two loci act epistatically to affect a trait under selection, change in one locus depends on, and tracks with, changes at the second locus (Takahasi and Tajima 2005). Empirically, Berg and Coop showed that the pattern of coordinated allele shifts for multiple complex traits such as height, skin pigmentation and T2DM differed from those under neutral evolution by comparing polygenic scores calculated from GWAS SNPs to the null distribution under random drift (Berg and Coop 2014), indicating the likelihood of such a model. Therefore, we posit that polygenic traits under selection should show evidence of coordinated allelic differentiation among populations, and that for two unlinked loci, variants should differentiate both associating loci simultaneously and with similar allele frequency changes. We modeled this possibility by extending the single locus ET method to a two-locus differentiation model, thereby accommodating potentially more complex genetic architectures.

As with single locus ET, by overlapping three interpopulation comparisons, we hypothesized that we would be able to substantially reduce background noise and enrich for two locus models that associate with complex phenotypes. If the underlying architecture includes more than a single locus, then two loci that coordinately differentiate among populations should associate with the same phenotypes more often than expected by chance. Such a framework should allow us to identify trait-associated SNP pairs, increasing the potential to detect putative epistasis and enrich our understanding of the genetic architecture of complex traits.

## Materials and methods

### HapMap 3 Genotype Data

The genotype data of individuals from four populations were downloaded from the International HapMap Project web site (Thorisson et al. 2005) (ftp://ftp.ncbi.nlm.nih.gov/hapmap/genotypes/latest_phaseIII_ncbi_b36/hapmap_format/consensus/). The selected samples were from Utah residents of Northern and Western European ancestry from the CEPH collection (CEU), Gujarati Indians in Houston, Texas (GIH), Yoruba in Ibadan, Nigeria (YRI) and African ancestry individuals from the Southwest USA (ASW) that had 1,440,504 SNPs in common (Supplementary Table S1). Mitochondrial and Y chromosomal SNPs were excluded due to their small number and different effective population sizes. We modified the overlapping pattern used in our analyses from that of the single locus ET method, where two high differentiation patterns (A vs. B and A vs. C, in Figure 1) and one low differentiation (B vs. C, in Figure 1) were used to identify the ET SNPs. In the two-locus model, low differentiation between populations with similar trait prevalence may obscure important patterns because it is necessary for risk alleles at both loci to be frequent for the associated phenotype to be prevalent. Under a two-locus model, low risk populations could have several allelic patterns: low risk allele frequencies at both loci; high risk allele frequency at one locus but low at the other. As a result, we assessed three different high comparisons among four populations (Figure 2, Venn diagram), CEU, YRI, GIH and ASW, assuming that CEU is divergent from the other three populations for a set of traits. We did not predefine any specific traits for these analyses as they were exploratory in nature and doing so could limit our ability to explore the full power of the two-locus method.

**Figure 2.**
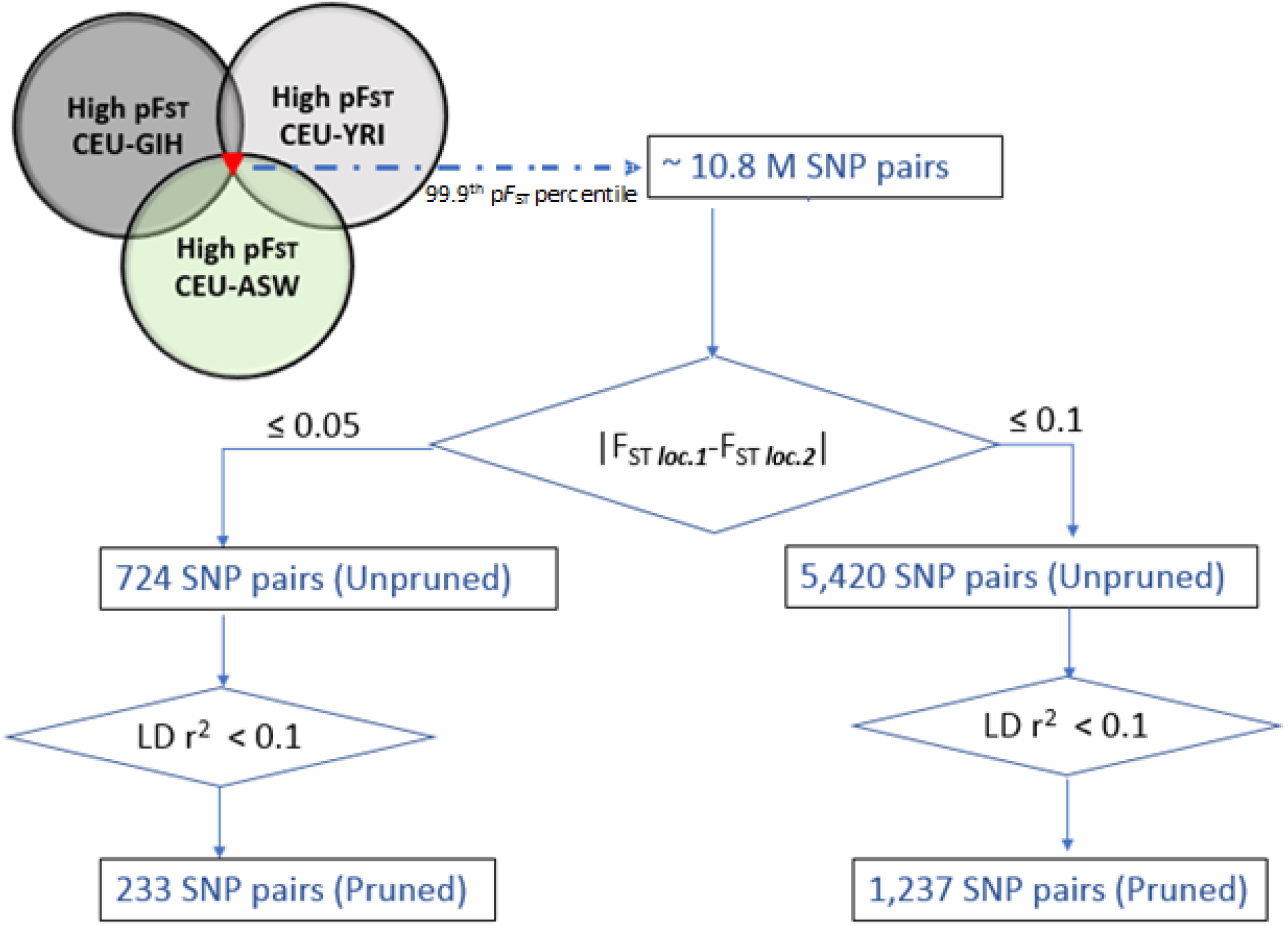
Flow chart representation of identification of co-differentiated ET SNP pairs among CEU, GIH, YRI and ASW populations. First, by applying 99.9^th^ p*F*_ST_ percentile thresholds, the number of overlapping SNP pairs reduced to approximately 10.8 million. Single locus *F*_ST_ difference thresholds of ≤ 0.05 or 0.1 in CEU vs. GIH, CEU vs. YRI and CEU vs. GIH were further applied to limit the dominance of p*F*_ST_ score by single locus differences, which reduced the number of ET SNP pairs to 724 and 5,420 respectively (two unpruned sets). Finally, LD pruning, with r^2^ < 0.1 threshold based on the CEU population, were applied to remove SNP pairs that were correlated with any other pairs at both loci, which generated 223 and 1,237 SNP pairs (two pruned sets).

### Estimating Two-locus Genetic Co-differentiation

Single locus *F*_ST_ was first calculated by a ratio of decomposed variance into three different components: *a* for between populations; *b* for between individuals within populations; and *c* for between gametes within individuals (Formula 1). The notations and calculations of *a*, *b* and *c* followed Weir and Cockerham (Weir and Cockerham 1984). The differentiation of each SNP pair (pairwise *F*_ST_, p*F*_ST_) was calculated by the weighted average over two loci (Formula 2), where *i* (= 1 or 2) denotes the *i*th locus.

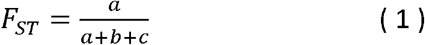

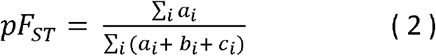

We then searched for overlap of highly differentiated SNP pairs, indicated by high p*F*_ST_ scores (99.9% percentile), between CEU and the other three populations YRI, GIH, and ASW (Figure 2, Venn diagram).

Since our aim was to search for co-differentiating SNP pairs, we excluded SNP pairs with a high p*F*_ST_ score due to a high single locus *F*_ST_ score, i.e. where one locus was highly differentiated while the differentiation on the second locus was small to moderate, which is unlikely in a two-locus epistatic system as noted above (Takahasi and Tajima 2005). Thus, we limited our search space to SNP pairs whose single locus *F*_ST_ value difference was ≤ 0.05 or ≤ 0.10, representing small to moderate thresholds of differentiation between the two loci, to control for extremely large single locus differences driving p*F*_ST_.

From each of the two *F*_ST_ difference thresholds, we obtained a set of SNP pairs based on the common HapMap SNP data across all four populations (the unpruned sets). To minimize the effects of correlation between SNP pairs within each *F*_ST_ difference threshold set, we applied a linkage disequilibrium (LD) threshold of *r*^2^ = 0.1 based on the CEU population to all ET identified SNP pairs. All SNPs in the same region were tested for LD, and only pairs where SNPs in both regions had less than the LD threshold were included, such that any SNP pairs from the same locus for which LD exceeded this value for both SNPs were removed from further consideration, thereby generating a pruned SNP pair set. The removal of SNPs in LD was random, in other words, one SNP was randomly chosen to represent a region if all SNPs in the region were of LD less than 0.1. In total, four sets of SNP pairs were assessed, two unpruned and two corresponding LD-pruned, from each *F*_ST_ difference threshold (Figure 2).

### Functional Annotation and Enrichment Analysis of Co-differentiated loci

For each set of ET SNP pairs, we mapped each SNP pair to gene pairs, using a 200 kb window around each SNP (±100 kb), so that one SNP pair could be mapped to multiple gene pairs. If each SNP in a pair mapped to a gene(s) associated with the same trait in GWAS catalog (Welter et al. 2014), we recorded the SNPs pairs and the trait. GWAS genes were defined by a p value threshold of 1 × 10^−6^. After scrutinizing all SNP pairs, we tallied the total number of co-associating SNP pairs for each trait. We then constructed a network from GWAS annotated pairs and traits using the network analysis software Gephi (Bastian et al. 2009). Nodes were defined as GWAS annotated genes and edges as shared traits. The degree of each node represents the number of edges emanating from each node and the weight of each edge represents the number of traits shared by each gene pair. We also assessed the structure of the network based on the degree distribution. If the distribution of edges per node follows a power law, then the network is scale-free and less likely to be random (Newman 2010).

To assess whether the SNP pairs from the ET filtering co-associated with a specific trait more frequently than expected by chance, a null distribution was generated by randomly selecting SNP pairs from the genome 10,000 times without matching allele frequencies, where each of the two SNPs were from different chromosomes. The number of selected SNP pairs for each random distribution was defined by the sizes of the unpruned and pruned sets. For the unpruned sets, random sets were produced regardless of the LD between each pair. For the pruned set, we selected SNP pairs such that no SNPs had r^2^ ≥ 0.1. Then, for each set of randomly generated SNP pairs, we tallied the total number of SNP pairs that have been associated with the same traits (Figure S1).

We determined the enrichment of ET-detected SNP pairs that co-associated with each specific trait, by comparing the number of SNP pairs from ET filtering to the null distribution generated from the 10,000 random SNP pair sets. The significance of the enrichment was indicated by an empirical *p*_*T*_ value for a trait *T* (Figure S1).

### SNP-SNP Interaction Effect of ET SNP Pairs in a Preterm Birth GWAS

We hypothesized that, for a given phenotype that differs in prevalence distribution among populations (e.g. the pattern in Figure 1), co-differentiated SNP pairs identified by ET are more likely to interact to affect the phenotype compared to SNP pairs identified as GWAS significant (index SNPs). To test this hypothesis, we asked whether index SNPs from a GWAS were more, or less, likely to exhibit epistasis than ET selected SNPs in the same chromosomal regions. We used existing preterm birth datasets to test this, based on its lower prevalence in CEU compared to GIH and YRI (Blencowe et al. 2012). These datasets contain genotype data of mothers from four different preterm birth cohorts that have previously been used to identify genetic associations of preterm birth: the Avon Longitudinal Study of Parents and Children cohort (ALSPAC) with 283 cases and 6,671 controls (Boyd et al. 2013; Fraser et al. 2013)(Supplemental Text); the Danish National Birth Cohort (DNBC) with 879 cases and 1,005 controls; the Finnish Birth Cohort (FIN) with 351 cases and 937 controls; the Genomic and Proteomic Network cohort (GPN) with 180 cases and 239 cohorts. For ALSPAC please note that the study website contains details of all the data that is available through a fully searchable data dictionary and variable search tool" and reference the following webpage: http://www.bristol.ac.uk/alspac/researchers/our-data/. Ethical approval for the study was obtained from the ALSPAC Ethics and Law Committee and the Local Research Ethics Committees. Informed consent for the use of data collected via questionnaires and clinics was obtained from participants following the recommendations of the ALSPAC Ethics and Law Committee at the time. For all cohorts only individuals of European ancestry were included in this analysis. Also, mothers with known risk factors for preterm birth or any medical complication during pregnancy influencing preterm birth, C-sections and non-spontaneous births were excluded. When assessing single locus associations in the maternal genome, six loci (*EBF1*, *EEFSEC*, *AGTR2*, *WNT4*, *TEKT3*, *KCNAB1 3’*) were significantly (5 × 10^−8^) associated with preterm birth as a dichotomous phenotype (Supplementary Table S2a), and six loci (*EBF1*, *WNT4*, *EEFSEC*, *AGTR2*, *ADCY5*, *KCNAB1 5’*) with gestational age as a continuous trait (Supplementary Table S2b) based on a meta-analysis between 23andMe GWA summary results (N=42,121) (Zhang et al. 2017) and these four data sets. As the two phenotypes were correlated, i.e. non-preterm birth status indicating longer gestational age, four loci (*EBF1*, *EEFSEC*, *AGTR2*, *WNT4*) were in both lists, but different index SNPs were identified that were in LD (*r*^2^ > 0.4) (Supplementary Table S2a, Table S2b and Supplementary Table S3). Based on the LD pattern between loci, SNPs with *r*^2^ greater than 0.4 were deemed as one locus in our analyses. The two *KCNAB1* SNPs were split into two loci as their LD was generally less than *r*^2^ of 0.4. We pooled the two lists of loci into one list of eight loci (*EBF1*, *WNT4*, *EEFSEC*, *AGTR2*, *ADCY5*, *TEKT3*, *KCNAB1 5’*, *KCNAB1 3’*). We then performed analyses on all pairs of these eight loci to generate a set of 28 or 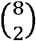 index SNP pairs. The interaction effects of each set of 28 index SNP pairs were then analyzed in regression models specified below in each of the four cohorts for the two phenotypes.

Given the high LD of index SNPs between the two GWAS lists, we also derived ET SNP pairs by merging loci from the two lists into regions based on LD structure. ET SNP pairs within ± 100 kb of the index SNP regions were identified using the following approach (Figure S2). Given the lower prevalence of preterm birth in CEU compared to GIH and YRI (Blencowe et al. 2012), we first chose ET SNP pairs based on their p*F*_ST_ scores between CEU and GIH/YRI, and between GIH and YRI. We used a score called Locus-Specific Branch Length (Shriver et al. 2004) (LSBL) to merge the three p*F*_ST_ scores into one value. Mathematically, the LSBL score measures the evolutionary distance (p*F*_ST_) of a SNP pair from CEU to both GIH and YRI together,

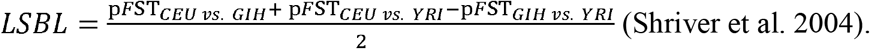

We calculated the LSBL scores of all inter-region SNP pairs between each two of the eight regions. Then SNP pairs with the highest LSBL scores were further filtered for single locus *F*_ST_ difference of less than 0.1 in CEU vs. YRI and CEU vs. GIH to prevent dominance by single locus *F*_ST_ signal, as above. If any SNP in an ET SNP pair was not found in the GWAS data (both genotyped and imputed), the pair containing this SNP was excluded from further analysis. We selected 100 ET SNP pairs with the highest LSBL scores to analyze their interaction effects in regression models specified below and compared with results to the 28 index SNP pairs. We modeled two types of outcomes: the dichotomous outcome, which is the status of preterm birth with preterm (≤ 37 weeks of gestational age) versus term birth (> 37 weeks of gestational age), and as a continuous outcome, i.e. gestational age. Regression analysis were run in the four different preterm birth cohorts, respectively.

For the dichotomous outcome, we tested the 28 index SNP pairs for SNP-SNP interaction effects using a logistic regression model with dichotomous preterm birth status as outcome, testing explicitly for the significance of interactions with no adjustments for covariates. We then assessed the significance of the SNP-SNP interaction between the 100 ET SNP pairs with the largest LSBL scores, using a logistic regression model with the dichotomous preterm birth status as outcome with no adjustments for covariates. For the continuous outcome, we tested the index and ET SNP pairs for epistatic effects using a linear regression model with gestational age as outcome in each cohort respectively, testing explicitly for the significance of interactions as above. We ran models with no covariate adjustments and also adjusted for maternal age and gender of infant. Finally, for all three models repectively, we compared the proportion of nominally significant interactions (p < 0.05) in both sets by two-tailed Fisher’s exact tests. The regression models can be expressed as below, where Y represents the outcome (dichotomous or continuous).

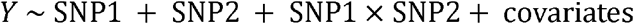

## Results and discussion

### Trait co-associations of ET co-differentiated SNP pairs compared random background

We calculated the unbiased two-locus p*F*_ST_ value for every inter-chromosomal SNP pair in the entire dataset, omitting all intra-chromosomal SNP pairs to exclude the possible effect of correlations between linked loci. This analysis generated approximately 10^12^ SNP pair comparisons (Figure 2; Supplementary Table S1).

Then we searched for the overlap of highly differentiated SNP pairs, indicated by extremely high p*F*_ST_ scores (99.9% percentile), among CEU vs. GIH, CEU vs. ASW and CEU vs. YRI (Figure 2, Venn diagram). The overlap consisted of 10,807,432 SNP pairs. We then excluded those pairs where p*F*_ST_ was dominated by single locus differences in all three comparisons between populations, using two thresholds, one where the absolute value of the difference in single locus p*F*_ST_ was ≤ 0.05 and one where it was ≤ 0.1. The former threshold reduced the number of SNP pairs to 724, and the latter to 5,420 pairs (Figure 2). We then queried the GWAS catalog to identify traits associated with genes in proximity to both ET SNPs in each pair. All ET-identified SNP pairs, mapped genes, associating traits are shown in the supplementary tables (Table S4 for the 724 set; Table S5 for the 5420).

For the 724 SNP pair sets, we identified two traits, schizophrenia and dental caries, where both ET SNPs were within ±100 kb of GWAS reported genes of that trait. Our method identified 12 ET SNP pairs for schizophrenia, an enrichment compared to chance alone based on results from randomly selected SNP pairs (p = 0.013). These were among the more than 100 loci that have been associated with schizophrenia (2011; Ripke et al. 2013). Consistent with our underlying hypothesis of differential distribution of ET identified diseases, individuals of African ancestry have substantially elevated rates of schizophrenia compared to European Americans after adjustment for socioeconomic status (Bresnahan et al. 2007).

Dental caries (DC) was also identified by our method. The role of genetics in DC has been clearly demonstrated by twin studies (Wang et al. 2010), and the etiology of DC involves interplay with environmental factors, such as diet. Our method successfully identified two SNP pairs associating with DC that showed a significant enrichment compared to randomly selected SNP pairs (p = 0.003). Ethnic disparities of DC exist among kindergarten students, after adjustment for poverty status, with white students having a significantly lower prevalence of DC compared to black students (Matsuo et al. 2015).

For the 5,420 SNP pairs set, 39 traits were identified using ET. Among these traits, 19 showed a significant enrichment in the identification of associated SNP pairs (Table 1). Twelve traits (denoted by an asterisk in Table 1) were skin or eye/hair color pigmentation related that are highly heritable and polygenic. Lighter skin pigmentation was clearly under selection in Europeans with selective pressures at high latitudes favoring lighter skin to facilitate vitamin D synthesis (Wilde et al. 2014). Another trait identified by our method, forced vital capacity (FVC, a lung function indicator), has also shown differences among populations. Caucasians have larger FVC compared with Asians and Africans (Whittaker et al. 2005; Menezes et al. 2015). To our knowledge, no significant difference has been reported in the five other traits (response to paliperidone, education attainment, obsessive-compulsive disorder, bipolar disorder, response to fenofibrate, Table 1) among the studied populations.

**Table 1.** Two-locus model of traits that were identified by ET in the unpruned set of 5,420 SNP pairs and the corresponding LD-pruned set of 1,237 SNP pairs. Empirical *p* value indicates the significance of ET enriches for two-locus models of these traits. Only traits where ET showing significant enrichment of two-locus models are shown. Traits marked with * indicates pigmentation related traits. NS indicates non-significant.

| **Trait** | **SNP pair count**<br>**in unpruned set**<br>**(*p* value)** | **SNP pair count**<br>**in pruned set (*p***<br>**value)** |
| --- | --- | --- |
| Basal cell carcinoma * | 5 (< 0.0001) | 3 (<0.0001) |
| Black vs. blond hair color * | 3 (< 0.0001) | 2 (<0.0001) |
| Black vs. red hair color * | 3 (< 0.0001) | 2 (<0.0001) |
| Eye color * | 3 (< 0.0001) | 2 (<0.0001) |
| Hair color * | 9 (< 0.0001) | 4 (<0.0001) |
| Perceived skin darkness * | 1 (< 0.0001) | 1 (<0.0001) |
| Skin color saturation * | 3 (< 0.001) | 2 (<0.0001) |
| Squamous cell carcinoma * | 5 (< 0.0001) | 3 (<0.0001) |
| Cutaneous squamous cell carcinoma * | 2 (0.00017) | 2 (0.0002) |
| Melanoma * | 2 (0.0112) | 2 (0.0003) |
| Obsessive-compulsive disorder | 4 (< 0.0001) | 1 (0.0008) |
| Skin sensitivity to sun * | 1 (0.0014) | 1 (0.001) |
| Tanning * | 1 (0.016) | 1 (0.0027) |
| Educational attainment | 9 (< 0.001) | 2 (0.0047) |
| Response to fenofibrate (HDL cholesterol levels) | 1 (0.032) | 1 (0.0082) |
| Lung function (FVC) | 4 (0.0019) | 1 (0.0161) |
| Immune response to smallpox (secreted IFN-alpha) | 1 (NS) | 1 (0.0185) |
| Response to paliperidone in schizophrenia (positive Marder score) | 59 (<0.0001) | 1 (0.0236) |
| Dental Caries | 7 (< 0.0001) | 1 (NS) |
| Bipolar Disorder | 10 (0.0025) | 1 (NS) |

To remove the dependence between the SNP pairs within each ET SNP set, two LD-pruned sets were also analyzed by the same method. Pruning was done by randomly selecting one SNP from those in high LD with each other at any given locus (r^2^ ≥ 0.1), which reduced the number of SNP pairs to 233 and 1,237 respectively (Figure 2). The results of random resampling were similar to those of the unpruned sets (Table 1). The GWAS reported SNPs, their effect sizes, and the LD between ET SNPs and are shown in Table S6 for the large pruned set with 1,237 ET SNP pairs.

### Network visualization of trait co-associations of ET co-differentiate loci

A network analysis of our results from the 5,420 ET identified SNP pairs was also constructed (Figure 3). Two genes were connected by an edge if they associated with the same phenotype. The degree distribution of our network follows a power law curve (*y* = 30.14*x*^−1.566^, R^2^ = 0.8804, Figure 3A), indicating that our network is unlikely to be random (Newman 2010). The network revealed more extensive co-differentiation than the random resampling results alone and it had 5 subnetworks (B to F). Subnetworks B and C each included several seemingly unrelated phenotypes, e.g. subnetwork C contained not only genes associated with all the pigmentation related traits but also with amyotrophic lateral sclerosis (ALS), obsessive-compulsive disorder, 3-hydroxy-1-methylpropylmercapturic acid levels in smokers, monobrow, immune response to smallpox, diisocyanate-induced asthma, response to fenofibrate and glomerular filtration rate (Figure 3). In subnetwork B, four genes associated with obesity-related traits (*GPHN*, *SDK1*, *MPHOSPH6* and *CTNNA3*) formed a simple connected structure, while the seven schizophrenia-associated genes formed a more complex cluster with interconnected nodes. The complex network structure of co-differentiated schizophrenia-associated genes may also reflect the more complex genetic architecture of schizophrenia. These network relationships indicate either that selection affects alleles associated with pigmentation acting together or that selection existed for some other phenotypes in the same subnetwork and the genes also associated with pigmentation. Based on prior evidence, the former is the more likely interpretation (Wilde et al. 2014). Both of these arguments depend on a highly pleiotropic genotype-phenotype map, affecting the differential rates of several traits simultaneously, which is likely (Chesmore et al. 2018).

**Figure 3.**
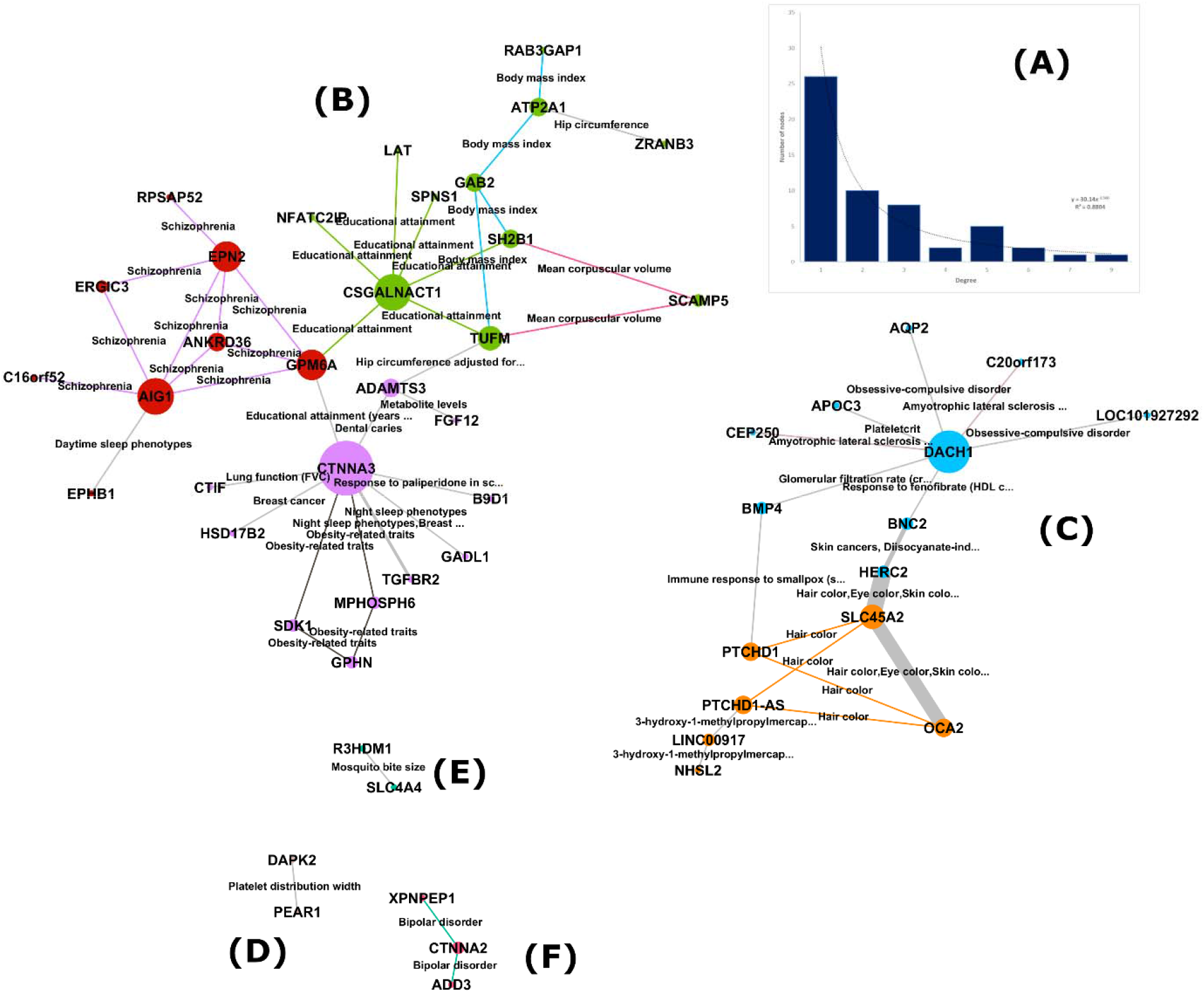
A network visualization of our results from the 5420 ET identified SNP pairs. (A) Degree distribution, showing scale-free like properties. (B-F) The network contains five different subnetworks). Nodes represent genes that were identified by ET, colored by their mathematically determined module. Node size reflects the number connections to other nodes. Edges represent shared phenotypes between gene pairs and thicker edges indicate multiple phenotypes. Edges are colored by phenotype.

Our two-locus model focused on loci that are physically unlinked, i.e. on different chromosomes. Independent assortment predicts that allele pairs from loci on different chromosomes will be randomized every generation and thereby inherited independently. Our analyses identified SNP pairs that we would not expect to differentiate simultaneously, but that did. Selection, therefore, in most cases is required to maintain these co-differentiations from unlinked loci. One possible scenario driving polygenic selection is epistasis. Under a two-locus epistatic disease model with unlinked loci, strong selection can drive different population prevalences. Two loci would be detected by our method only if they had both contributed to the trait under selection and did so under a model where both varied in concert. For example, skin pigmentation, a polygenic trait, is clearly under strong selection (Wilde et al. 2014). Given that 12 skin pigmentation related traits were found using pairwise ET (Table 1), it is clear that our method works for traits that are strongly selected. Though the evidence for selection may be weaker for other traits, we still observed evidence of enrichment for coordinated allele frequency changes among the studied populations in traits that vary in prevalence, such as schizophrenia, dental caries and lung function. Our results support two locus models for 19 out of 39 traits (18 out of 38 for the pruned set) identified in our analyses, a substantial and significant enhancement over random expectations.

In the study design, we filtered for SNP pairs that had small to moderate differences in single locus *F*_ST_ measures (*F*_ST_ ≤ 0.05 or 0.1) with the intent of only identifying co-adapted SNP that exhibited a coordinated shift of allele frequencies. This approach prevents focusing on SNP pairs where the p*F*_ST_ is dominated by large patterns of differentiation at a single locus, as explained in the methods section. This should enhance the ability to detect epistasis as opposed to two loci acting additively because coordinated changes can only be maintained by genes acting together or through interactions when they are unlinked. For example, if two SNPs affect the prevalence of a trait, and do so additively, selection on the trait should change their allele frequencies independently and primarily as a function of their relative effect sizes. In contrast, if the effects of these two SNPs are primarily epistatic, their allele frequencies should change in concert; in other words, changes in allele frequency at one locus of an epistatic pair would depend on the changes at the second locus thereby creating a coordinated pattern of allele frequency change at both loci. This will be detectable in p*F*_ST_ when the two single-locus *F*_ST_ values are similar. Thus, our method has the potential to reveal putative epistasis in the genetic architecture of complex phenotypes.

Compared to single locus ET, the two-locus method was able to identify genes associated with more complex diseases, such as schizophrenia, which is notable for having a large amount of missing heritability (van Dongen and Boomsma 2013). This is likely due to our extension of the ET method that can identify multi-locus effects, and thereby a more complex underlying genetic architecture, such as those that incorporate epistasis implicitly. Epistasis is also frequently suggested as a potential explanation for the missing heritability observed in genome-wide association studies, although this hypothesis still has a very limited evidentiary basis (Zuk et al. 2012); our findings provide additional supportive evidence for a more complex architecture of the traits we identified.

As evidence that our method can detect epistasis, we found the genes *HERC2* (rs12913832) and *SLC45A2* (rs16891982) that interact in Europeans to affect eye color (Wollstein et al. 2017). It has been argued, though, that *HERC2* is actually tagging *OCA2*, a known eye-color gene (Eiberg et al. 2008). The coordinated allelic differences among populations and their correlation with known two locus interactions that appear to exist serves as a proof of principle that our analyses of p*F*_ST_ can reveal underlying epistasis. Similar patterns have been observed before. For example, two interacting immune response loci *HLA* and *KIR*, although on different chromosomes, showed significant correlation in their allele frequencies among different populations, likely driven by selection pressure from pathogens (Single et al. 2007). Additionally, introgressed regions between Neanderthals and modern humans showed significantly lower divergence from patterns of variation in the genomic background; this can be explained by the association between high divergence and negative epistasis that leads to allelic incompatibility (Vernot and Akey 2014). In the study of a non-human system, swordtail fish, introgressed regions showed higher divergence, which could be explained by more, but weaker, purifying selection or that these regions are less functionally essential (Schumer et al. 2016).

It is worth noting that our empirically derived p values depend on the number of loci in the GWAS catalog. For a trait that is highly polygenic, e.g. schizophrenia, with more than 100 reported associating variants in the GWAS catalog, we would expect the randomly sampled SNP pairs to be from associating genes more often than a less complex trait. This will essentially reduce evidence for significant enrichment using p*F*_ST_, as we found. Also, our estimation of genetic effects of SNPs on phenotypes are based on GWAS results, which are theoretically based on estimating the additive genetic variance of a trait by decomposing the total variance into each SNP in an additive manner. However, additive variance does not necessarily reflect biologically additive gene actions in a phenotype; neither does the existence of additive variance indicate that genes act additively. Detected additive variance can be a mix of both additive and epistatic effects (Cheverud and Routman 1995; Maki-Tanila and Hill 2014). Therefore, our two-locus ET method can be used to further dissect the underlying architectures of seemingly marginal associations, which will help better elucidate true genetic models.

### Selecting epistatic models using two-locus ET results from studies of Preterm Birth

Our two-locus ET method showed the potential ability to detect co-differentiation at two loci driven by epistatic selection. Given that a trait has different prevalences among populations, we proposed that co-differentiated SNP pairs identified by ET were more likely to interact to affect the phenotype compared to GWAS identified SNPs (index SNPs). We tested this hypothesis using preterm birth GWAS datasets. Six loci (*EBF1*, *EEFSEC*, *AGTR2*, *WNT4*, *TEKT3*, *KCNAB1 3’*) were previously associated with preterm birth, and six loci (*EBF1*, *WNT4*, *EEFSEC*, *AGTR2*, *ADCY5*, *KCNAB1 5’*) with gestational age (Table S2). As the two phenotypes were highly correlated, multiple index SNPs in the two sets were in high LD (Table S3). As we discussed in our method section, if two SNPs had LD *r*^2^ greater than 0.4, we considered them interchangeable in the regression analyses. Together there were eight index loci (*EBF1*, *EEFSEC*, *AGTR2*, *WNT4*, *TEKT3*, *KCNAB1 3’, ADCY5, KCNAB1 5’*), which corresponded to 28 index pairs and thus 112 tests total for epistasis across the four cohorts (4 × 28). For ET SNP pairs, the total number of epistasis tests is 400 across the four cohorts (4 × 100 candidate SNP pairs based on LSBL scores).

In the analysis using preterm birth status as a dichotomous outcome, three index SNP pairs showed significant interaction effects, but none of the significant interactions replicated across the cohorts. In contrast, 31 ET identified two-locus models had significant interaction effects, and six SNP pairs were replicated in two cohorts (Table 2a). The SNP pair replications represented two gene pairs, *ADCY5* - *KCNAB1* 5’ and *EEFSEC* - *KCNAB1* 3’. ET SNPs identified slightly higher proportion of two-locus models compared to the index SNPs; however, it was not statistically significant (Fisher’s exact test *p* = 0.083). Nonetheless, ET SNP pairs exhibited robust replication across multiple cohorts compared to no replication in the paired index loci. Interacting loci identified by ET and index SNPs were all different providing a novel set of putative epistatic loci (Table 2a, Figure 4a).

**Table 2a.**
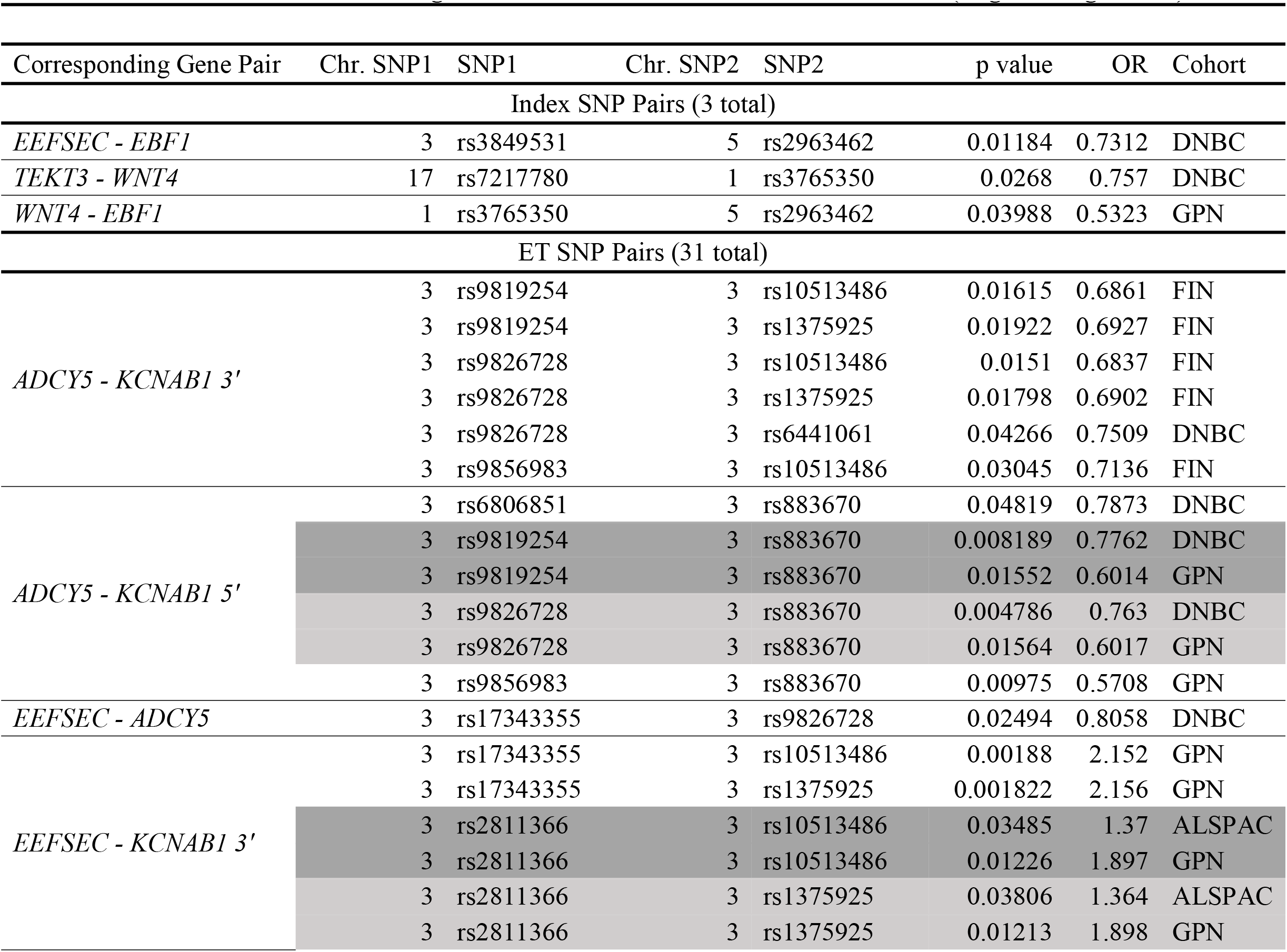
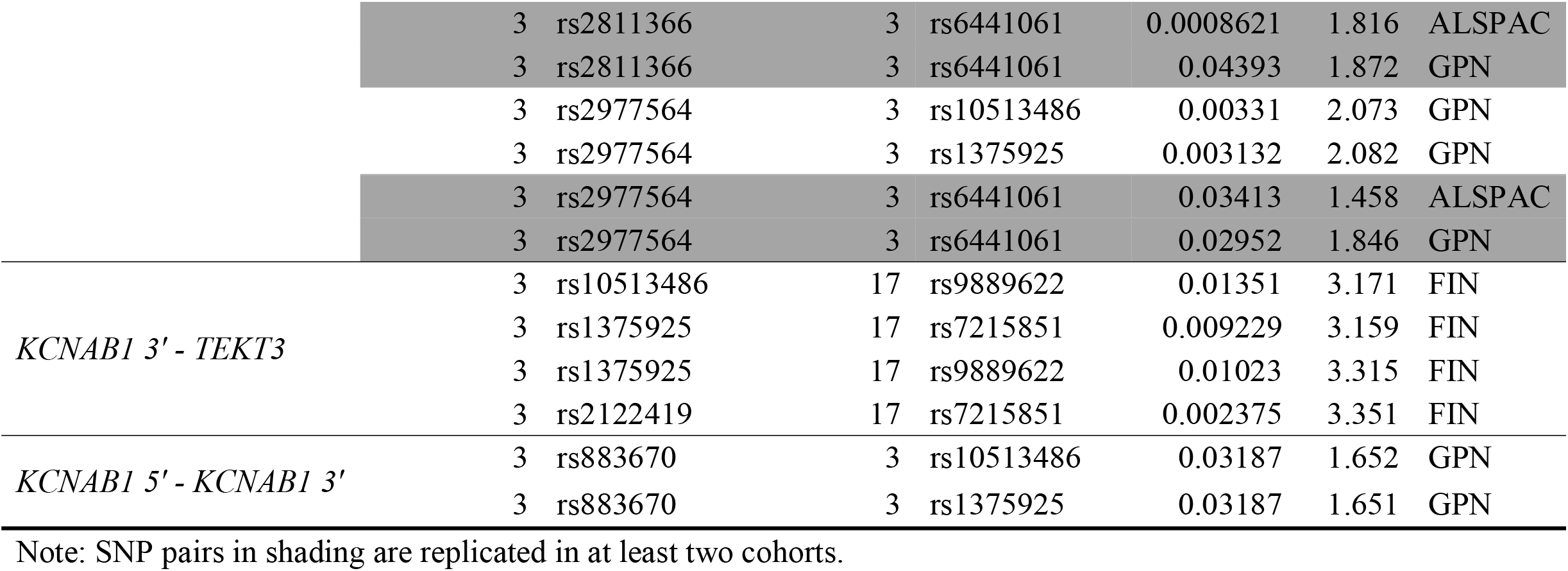
SNP Pairs with a Significant Interaction Effect for Preterm Birth (Logistic Regression)

**Figure 4.**
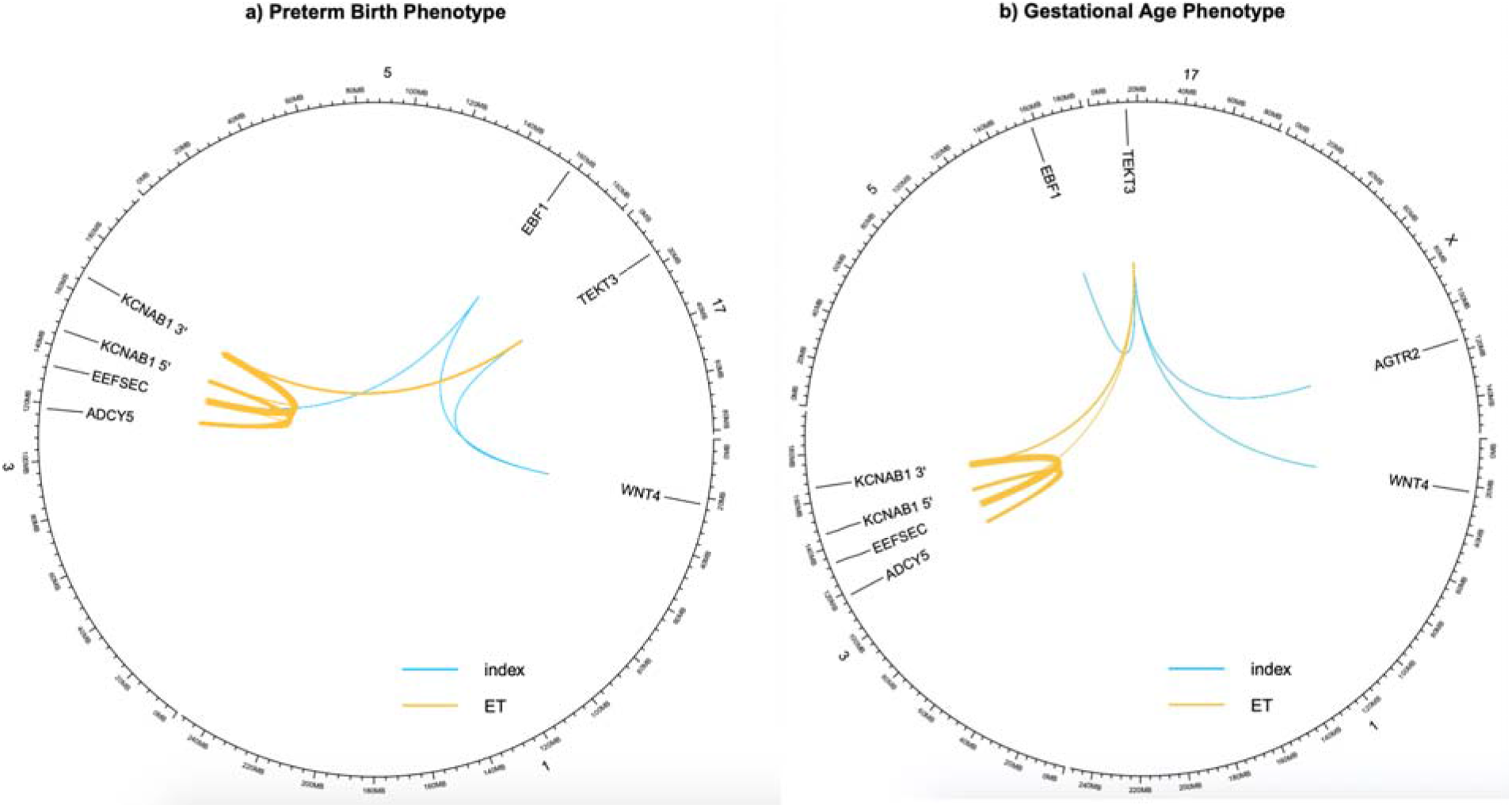
Two-locus epistasis models of preterm birth status or gestational age. A circus plot of epistatic loci in the genome. Two loci are linked if they show a significant interaction effect on the preterm birth status (a) or gestational age (b, model with covariates) in regression analyses. The thickness of a link is proportional to the number of interacting SNPs pairs identified, including replication across cohorts. The coordinates on chromosomes are not exact positions of loci.

In the linear regression analysis, we first modeled gestational age as the outcome with no adjustments for any covariates in the same four cohorts. In total, 34 ET SNP pairs showed significant interaction effects, while only 3 index SNP pairs showed significant interaction effects (Table 2b). Under this regression model, ET identified more SNP interaction models compared to index SNP pairs (Fisher’s exact test *p* = 0.038). However, when we adjusted for maternal age and infant sex in the model, the number of significant interactions in the ET and index SNP pairs became 29 and 3 (Table 2c). The difference in the proportion of epistatic interactions between the ET and index SNP pairs was not significant (Fisher’s exact test *p* = 0.081), but again there were new gene pairs that provided evidence for significant interaction (Figure 4b). Similar to the results from the logistic regression, none of the index SNP replicated across cohorts, but seven ET SNP pairs replicated in at least two cohorts and two pair replicated across three cohorts (Table 2c). The interacting gene pairs replicated were *ADCY5* - *KCNAB1* 5’ and *EEFSEC* - *KCNAB1* 3’, the same ones as for preterm birth as a dichotomous trait (Table 2c).

**Table 2b.**
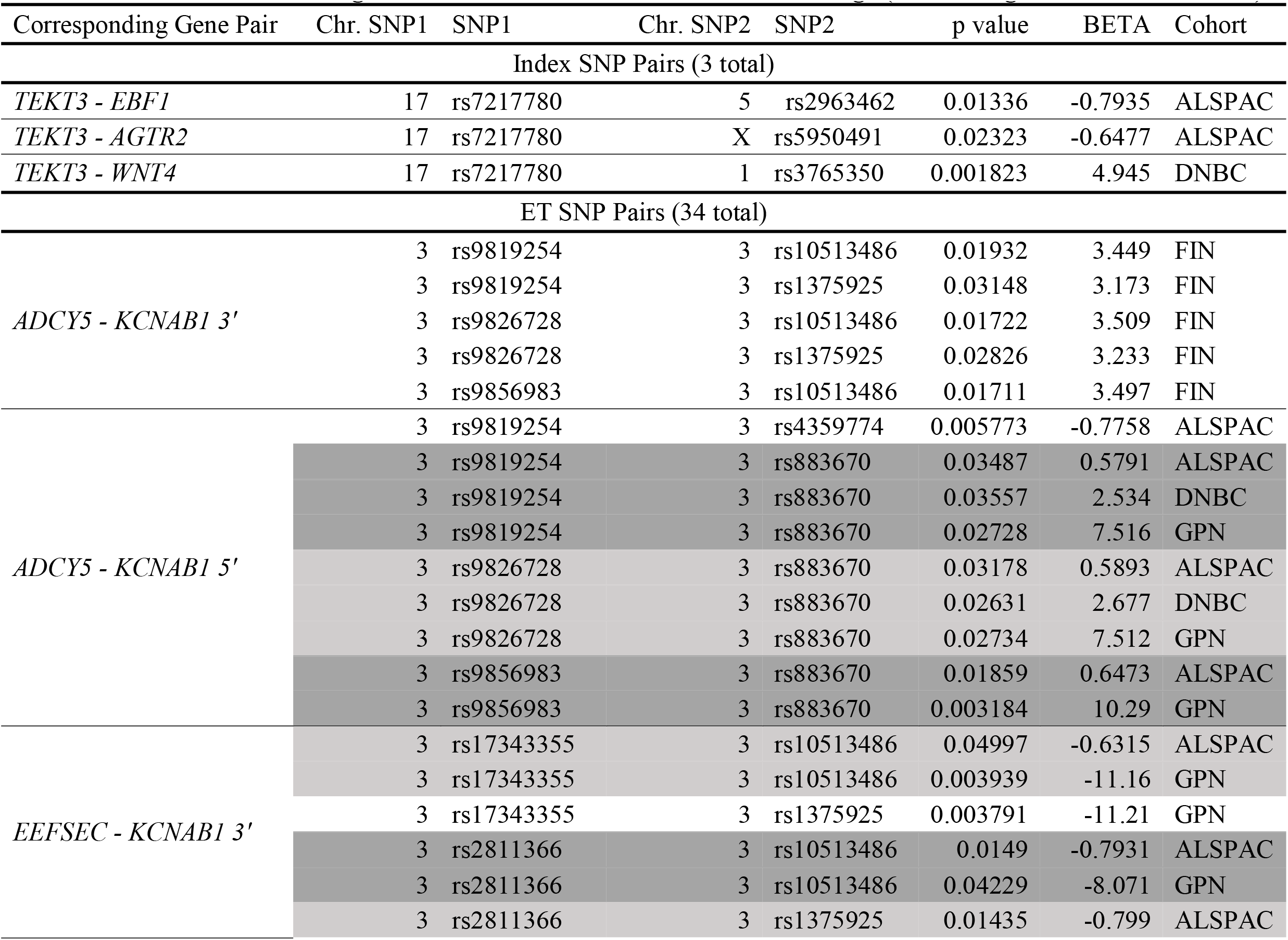
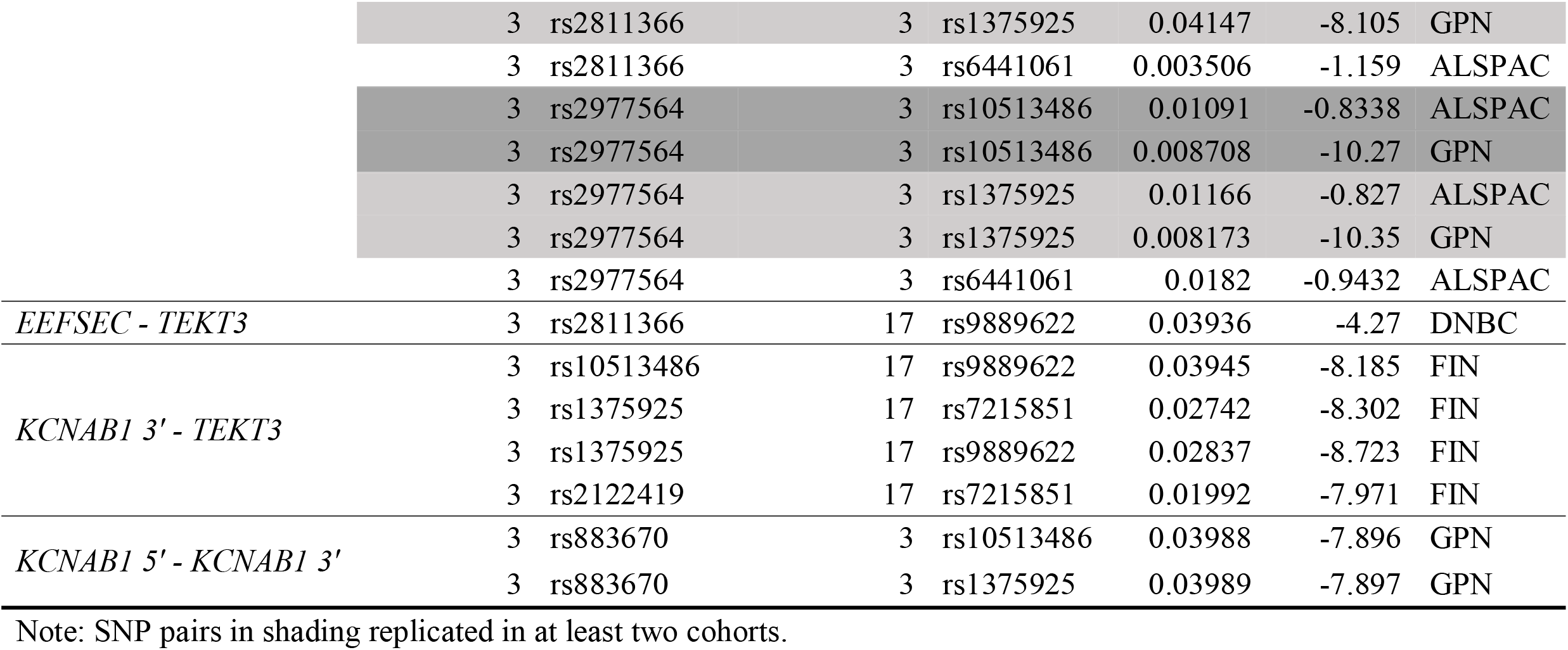
SNP Pairs with a Significant Interaction Effect for Gestational Age (Linear Regression, no covariates)

**Table 2c.**
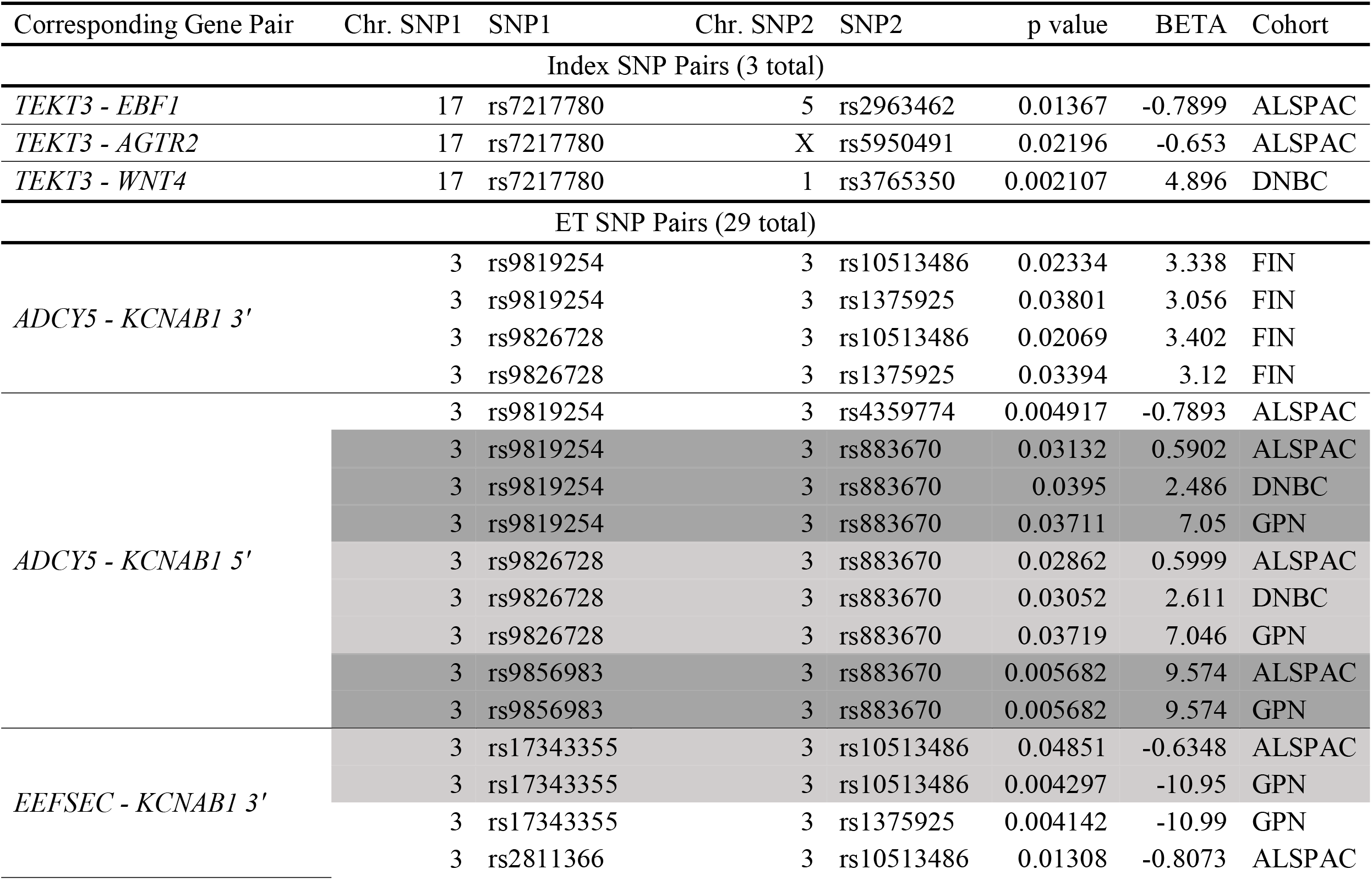
SNP Pairs with a Significant Interaction Effect (Linear Regression, with covariates)

Several notable additional results were generated using ET to detect epistasis among candidate loci for pregnancy outcomes generated from the GWAS analyses. ET SNPs not only identified at least the same proportion of epistatic models compared to the index SNPs, but also, they identified putative interactions among loci that the index SNPs did not. For example, an interaction model was evident between *ADCY5* and *KCNAB1 5’*. ET SNPs showed significant interactions in both logistic regression models and the linear regression models (Table 2a,2b and 2c, Figure 4). More importantly, ET identified putative interactions that were replicated across multiple cohorts and thus more likely to be real interactions. For instance, the interaction between *ADCY5* and *KCNAB1 5’* was identified by ET in three cohorts in the linear regression models and in two cohorts in the logistic regression model (Table 2a,2b and 2c). In contrast, index SNPs did not replicate any interaction models across cohorts nor did they identify any two-locus models between *ADCY5* and *KCNAB1 5’* for preterm birth status (logistic regression) or gestational age (linear regression). These two genes show no signs of interaction in the databases that we screened, but on assessing putative biological interactions, ADCY5 is involved in cyclic adenosine monophosphate (cAMP) production (Ding et al. 2004) and many potassium channels are cAMP dependent (e.g. KCNQ1 and KCNE1)(Kurokawa et al. 2003), although supporting data for this function for KCNAB1 has not been found yet. In vitro experiments in both human and mouse myometrial tissues have shown the possibility of potassium channels as drug targets of preterm labor; a potassium channel inhibitor could significantly increase spontaneous myometrial contractions while potassium channel activators could lead to myometrial relaxation (McCallum et al. 2011).

Taken together, our results indicate that there are likely real and robust findings for two locus models affecting pregnancy outcomes, discovered agnostically with respect to both function and index SNP. Other novel interactions identified by ET but not with index SNP interaction models (in at least one regression analysis, linear or logistic), include *KCNAB1 3’* - *EEFSEC*, *TEKT3* – *EEFSEC*, *TEKT3* - *KCNAB1 3’*, *KCNAB1 3’* - *ADCY5* and *KCNAB1 3*’ - *KCNAB1 5’* (Table 2a, 2b and 2c, Figure 4). These results show that ET filtering can identify SNP-SNP interactions for preterm birth, a highly complex trait, not evident with interaction analyses based solely on index SNPs. Therefore, ET filtering as a method provides additional epistatic candidates for further study. These results, especially the case in which epistasis replicated (*ADCY5* - *KCNAB1 5’*), indicate that ET can detect novel interactions that are as, or more, likely to be real than index SNP interactions. Of course, these new putative interactions will require explicit and independent validation, but the results indicate that Index SNP analyses alone is insufficient when considering the possibility of epistasis.

## Conclusion

We found that polygenic effects for multiple complex traits, including pigmentation related traits, associated with several pairs of loci that evolved in concert, an unlikely scenario if the two locus effects were purely additive. In addition, we showed that SNP pairs defined by ET characterized novel interactions that replicated across several cohorts for preterm birth and gestational age. Index SNPs for these same traits were not as robust. It is probable that co-differentiating SNP pairs detected by our method are influencing multiple complex traits in an epistatic manner, and our method can refine the genetic architecture of complex traits due to epistatic interactions. In conclusion, by using co-differentiation of unlinked loci among multiple populations, we were able to identify several traits for which epistasis is a more likely model than simple additive effects among loci.

## Acknowledgements

We are extremely grateful to all the families who participated in Avon Longitudinal Study of Parents And Children (ALSPAC), Finnish Birth Cohort (FIN), Danish Birth Cohort (DNBC) and Genomic and Proteomic Network (GPN). Our sincere thanks to dbGaP for depositing and hosting data access for the current research. The DNBC datasets used for the analyses described in this manuscript were obtained from dbGaP at http://www.ncbi.nlm.nih.gov/sites/entrez?db=gap through dbGaP accession number phs000103.v1.p1. The GPN genetic data was acquired from dbGaP at Accession phs000714.v1.p1

This work is supported by a grant from the March of Dimes to the MOD Prematurity Research Center Ohio Collaborative and grants LM010098 and AI116794 from the National Institutes of Health. Genetic data for the ALSPAC was supported by the UK Medical Research Council and Wellcome (Grant ref: 102215/2/13/2) and University of Bristol provide core support for ALSPAC. This publication is the work of the authors and Minjun Huang and Scott Williams will serve as guarantors for the contents of this paper.

